# An automated approach to quantify chemotaxis index in *Caenorhabditis* nematodes

**DOI:** 10.1101/2022.04.30.490142

**Authors:** Timothy A. Crombie, Chido Chikuturudzi, Daniel E. Cook, Erik C. Andersen

## Abstract

Chemotaxis assays are used extensively to study behavioral responses of *Caenorhabditis* nematodes to environmental cues. These assays result in a chemotaxis index (CI) that denotes the behavioral response of a population of nematodes to a particular compound and can range from 1 (maximum attraction) to −1 (maximum avoidance). Traditional chemotaxis assays have low throughput because researchers must manually setup experimental populations and score CIs. Here, we describe an automated methodology that increases throughput by using liquid-handling robots to setup experimental populations and a custom image analysis package, ct, to automate the scoring of CIs from plate images.

## Description

The *Caenorhabditis elegans* chemosensory system can detect a variety of volatile (olfactory) and water-soluble (gustatory) compounds associated with nutrients, danger, or competitors in their natural environment (Cornelia I. Bargmann 2006). In response to some chemicals, nematodes will move towards the cue (chemotaxis) or actively avoid it by moving away (Cornelia I. Bargmann 2006). Behavioral responses to environmental cues are often studied using a chemotaxis assay, where the movement of animals on an assay plate is recorded after an exposure to a compound. These assays yield a chemotaxis index (CI) that denotes the behavioral response to the particular compound on a scale ranging from 1 to −1, *i.e*., maximum attraction to maximum avoidance, respectively. Over the last 50 years, various chemotaxis assay protocols have been used to study the genes, molecular mechanisms, and neuronal circuits underlying olfaction, olfactory preferences, and the control of chemotactic behaviors in *C. elegans* (Ward 1973; C. I. Bargmann, Hartwieg, and Horvitz 1993; Troemel, Kimmel, and Bargmann 1997; Itskovits et al. 2018). These assays have also been central to the discovery that *C. elegans* can learn to adapt to olfactory cues through the formation of short and long-term memories (Kauffman et al. 2011; Colbert and Bargmann 1995), and that *C. elegans* demonstrate innate preferences for bacterial food sources (Zhang, Lu, and Bargmann 2005). However, most of this research was performed using a single laboratory-adapted genetic background, N2, which means that relatively little is known about how natural genetic variation influences these traits. The few studies exploring natural genetic variation have revealed that different *C. elegans* strains have distinct preferences among bacterial species and vary in their ability to distinguish among bacterial food sources (Glater, Rockman, and Bargmann 2014; Volkers et al. 2013). One of the largest challenges to exploring the role of natural genetic variation in behaviors driven by the chemosensory system is the limited throughput of traditional chemotaxis assays because they require manual setup and scoring.

One well established chemotaxis assay protocol measures CIs using standard 6 cm NGM plates divided into quadrants as a test arena (Margie, Palmer, and Chin-Sang 2013). In this design, developmentally synchronized nematodes are transferred to the center of a plate (origin), a test compound is spotted in two of the four quadrants, and a control compound is spotted in the other quadrants, all spots are equidistant from the origin and are mixed with sodium azide to paralyze animals once they reach the spot. After the nematodes explore the plate for one hour, the number of animals in each quadrant are counted and the CI is calculated as the number of animals in the test quadrants minus the number of animals in the control quadrants divided by the total number of animals that have left the origin. Although this methodology is simple to use and provides robust results, it is difficult to scale to large experiments because it requires researchers to manually observe and record the position of every nematode on each plate.

Here, we describe a chemotaxis assay methodology that increases throughput by using liquid-handling robots to setup chemotaxis plates and custom image analysis scripts to automate the calculation of CIs from plate images. Our protocol begins by propagating strains for three generations to control for the transgenerational effects of starvation on the source plates (Figure 1-A1). Gravid adults are bleach synchronized to collect embryos then the embryos are titered in K medium to 1 embryo/μL and left overnight without food to arrest as L1s. The following day, approximately 1000 arrested L1s are transferred to each of three 6 cm NGMA plates per strain and grown at 20°C for 48 hours until they reach the L4 larval stage (Figure 1-A2). We wash the L4s off the plates with M9 and use the COPAS BIOSORT (Union Biometrica; Holliston, MA) to dispense 50 developmentally synchronized fourth larval stage (L4) animals to the origin of 6 cm chemotaxis plates. To accurately dispense nematodes, we fabricated a custom plate holder from acrylic that fits securely on the BIOSORT stage and accommodates six chemotaxis plates (Figure 1-A3). After nematodes are dispensed to the six plates, the plate holder is removed from the BIOSORT stage and test compounds are added to the quadrants using a multi-channel pipette. The plate holder is designed so that multiple test compounds can be added in a single pipette step. The exact positioning of the test spots is guided by targets laser etched into the plate holder (Figure 1-A4). Once the compounds are added, the plates are removed from the holder and transferred to a 20°C incubator for one hour to allow the animals to respond to the test compounds (Figure 1-A5). Afterwards, the plates are transferred to 4°C to arrest animal movement and await imaging. Working in tandem, a pair of researchers can quickly process chemotaxis plates by using two plate holders and cycling through the assay steps six plates at a time. The plates can be left at 4°C for up to five days prior to imaging. Any microscope with a camera attachment and an appropriate field of view can be used to acquire images, but we used a previously described imaging system (Churgin and Fang-Yen 2015).

**Figure 1.**
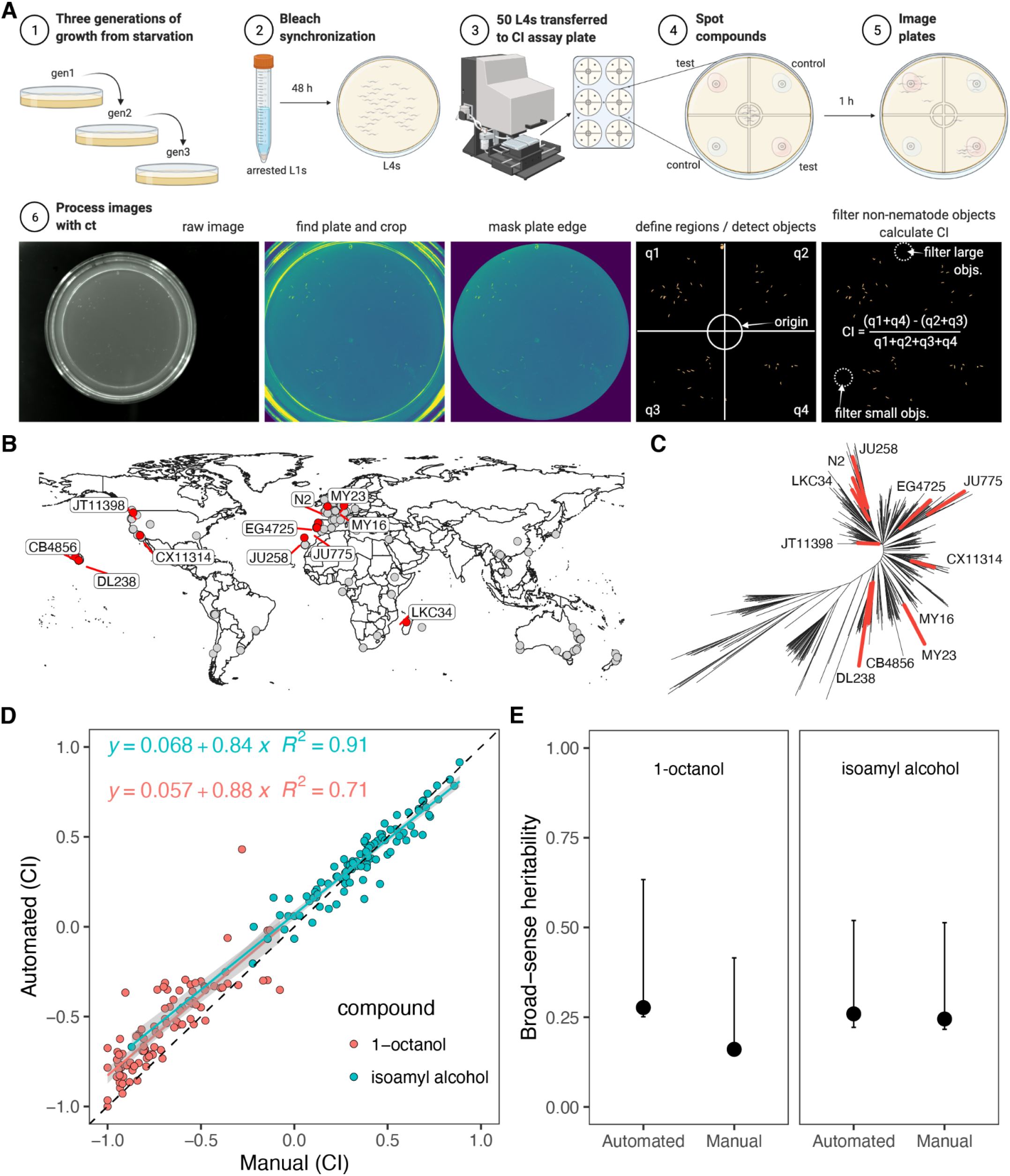
An automated assay workflow yields robust chemotaxis measurements. **(A)** The experimental workflow is shown. **(A1-3)** Nematodes are grown for three generations prior to being transferred to chemotaxis assay plates as synchronized fourth larval stage animals (L4s). **(A4)** Compounds are added to the chemotaxis assay plates using the guides (gray) that are laser etched into the custom acrylic plate holder and are visible through the NGMA plates. **(A5)** The animals are left to respond to cues for one hour at 20°C then the plates are transferred to 4°C before imaging. **(A6)** The image processing steps performed by ct are shown. The steps result in an output file with the chemotaxis index (CI) calculated for each image passed to the ct command. **(B)** A map of isolation locations for the 11 strains used in this experiment (red points), the other wild *C. elegans* strains available from the *Caenorhabditis elegans* Natural Diversity Resource (CeNDR) are also shown (gray points) (release 20210121) (Cook et al. 2017). **(C)** The genetic relatedness of CeNDR strains are shown as an unrooted tree, and the 11 strains used here are colored red and the other 529 CeNDR strains are colored black. **(D)** The relationship between manual CI calculations and automated CI calculations for identical images is shown. The coefficient of determination (R^2^) and the linear regression equation are shown for each compound. The linear models were calculated using CIs of all 11 strains for either 1-octanol or isoamyl alcohol. **(E)** Estimates of broad-sense heritability using data from all 11 strains are shown for each compound using either the manual or automated CIs. The black dots are point estimates, and the error bars represent 95% confidence intervals calculated by bootstrapping.

In our assay, the quantification of CIs from plate images is automated using our custom command line utility ct, which is written in Python and easily installed under Linux/OSX systems using the instructions at the following GitHub repository: https://github.com/AndersenLab/chemotaxis-cli. ct identifies the plate boundaries in a raw image, segments objects on the plate from the background, filters out non-nematode objects, assigns quadrant positions to the nematode objects on the plate, measures the total area of nematode objects within each quadrant, then calculates a CI for a particular plate without the need for manual counting (Figure 1-A6). Importantly, ct does not attempt to count individual nematodes because this leads to errors when nematodes are clumped together tightly on plates (Wählby et al. 2012), instead the pixel area of nematode-objects is used as a proxy for nematode counts. ct outputs intermediate image files for debugging purposes, and a results file in text format with rows for each image processed and several data fields for each row, including the number of pixels for nematode objects in each quadrant, the total number of non-nematode pixels filtered from the image, and the CI for the image. Detailed installation and usage notes for ct are available at the GitHub repository linked above.

We validated our automated approach by comparing the CIs calculated with ct to CIs calculated using manual counts of nematodes in the same images. For this test, we measured CIs to a known repellant, 1-octanal, and a known attractant, isoamyl alcohol (Chao et al. 2004; C. I. Bargmann, Hartwieg, and Horvitz 1993). To test whether ct would work when applied to different genetic backgrounds, we also performed the comparisons across 11 different wild strains isolated from around the globe (Figure 1-B,C). We found that the two methods were highly correlated (*R^2^* = 0.91) when comparing CIs for isoamyl alcohol across 11 different *C. elegans* strains (Figure 1-D). The methods were less correlated when comparing CIs for 1-octanol (*R^2^* = 0.71) but still highly correlated. We believe one the reason that ct performed worse for 1-octanol relative to isoamyl alcohol is because 1-octanol is weakly soluble in water and the edge of the unabsorbed test spot was occasionally detected as a nematode object by ct. This limitation could be overcome by careful adjustment of the plate illumination so that unabsorbed test spots do not reflect illumination light in the images. We calculated the broad-sense heritability (*H^2^*) of CIs among our strains using either the automated or manual CI estimates (Figure 1-E). The *H^2^* estimates indicate the proportion of total phenotypic variance explained by strain differences and are useful metrics to assess whether genetic variation influences a trait of interest. We found that *H^2^* estimates were within a similar range when calculated from either the automated or manual CIs, suggesting that ct could be useful for dissecting the natural genetic variants underlying differences in chemotaxtic behavior among wild *C. elegans* strains.

This study describes a new methodology to increase the throughput of chemotaxis assays while maintaining assay accuracy. We propose that this method could be used to facilitate large-scale genetic mappings of behavioral responses to various test compounds.

## Methods

### Strains

Animals were grown at 20°C and fed *E. coli* OP50 grown on modified nematode growth media (NGMA) containing 1% agar and 0.7% agarose to prevent burrowing (Andersen et al. 2014). We used the strains CB4856, CX11314, DL238, EG4725, JT11398, JU258, JU775, LKC34, MY16, MY23, and N2. All strains are available from the *C. elegans* Natural Diversity Resource (CeNDR) (Cook et al. 2017).

### Chemotaxis Assay

Prior to the start of the assay, the strains were each transferred for three generations then bleached to collect embryos. The embryos were then titered in K medium to 1 embryo/μL and left overnight without food to arrest as L1s. The following day, approximately 1000 L1s were transferred to each of three 6 cm NGMA plates per strain. Then, the animals were grown at 20°C for 48 hours, washed from the plates with M9, and sorted as L4s onto 6 cm chemotaxis assay plates using the COPAS BIOSORT (Union Biometrica; Holliston, MA) and a custom fabricated plate holder: https://github.com/AndersenLab/chemotaxis-cli/tree/master/customPlateHolder. The test compounds or experimental controls were then spotted on the plates using the holder guides for placement. All spotting solutions were prepared with 1 M sodium azide to paralyze animals as they encountered the spots (Margie, Palmer, and Chin-Sang 2013). After one hour at 20°C, the plates were transferred to 4°C and imaged within 72 hours. The chemotaxis assays were performed in three experimental blocks. Each block contained all 11 strains with three replicate chemotaxis plates per strain per test compound.

### Statistical analysis

All statistics and plotting were performed using R statistical software (v4.1.1) (R Core Team 2020). Linear models were fit to CI data to compare assay methods using the *lm* function in the stats package. To calculate broad-sense heritability *(H^2^*), we first used the *lmer* function in the lme4 package to model the CI phenotypes with strain as a random effect (*phenotype ~1 + (1|strain)*) (Bates et al. 2015). We then calculated *H^2^* as the fraction of the total phenotypic variance that could be explained by the random component (strain) of the mixed model as described previously (Zdraljevic et al. 2019).

## Author Contributions

Chido Chikuturudzi: Conceptualization, Investigation, Methodology, Visualization, Writing – review & editing. Timothy A. Crombie: Conceptualization, Investigation, Methodology, Software, Formal Analysis, Visualization, Writing – original draft, Writing – review & editing. Daniel E. Cook: Conceptualization, Methodology, Software. Erik C. Andersen: Conceptualization, Supervision, Writing - review and editing.

## Acknowledgments

We would like to thank Ezequiel Alvarez-Saavedra for the design and fabrication of the chemotaxis plate holder we used with the COPAS BIOSORT. We would also like to thank members of the Andersen laboratory for their helpful comments on the manuscript.

